## Supplementary Information for "Contractility-induced self-organization of smooth muscle cells: from multilayer cell sheets to dynamic three-dimensional clusters"

Wang *et al.*

### Supplementary Information

- Supplementary Figures
- Supplementary Notes
- Supplementary Movies

### Supplementary Figures

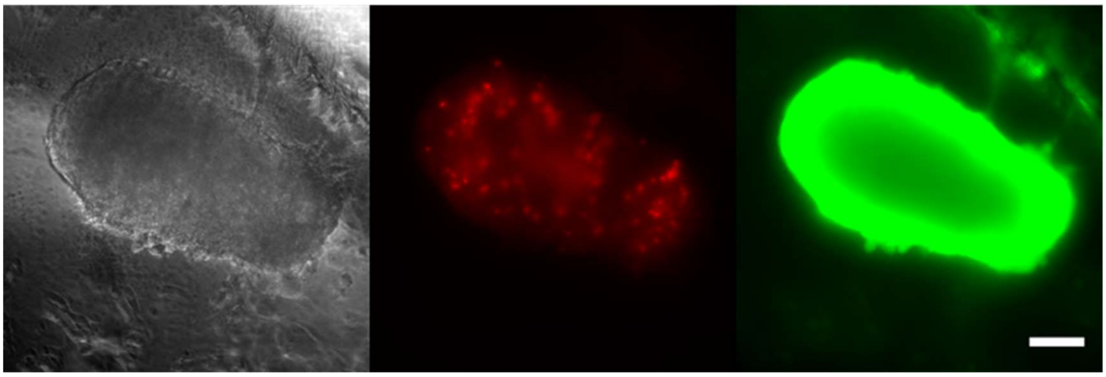

**Figure S1.** Left: Brightfield image of a SMC cluster. Right: Live-dead staining of SMCs in a cluster. Red: dead cells; green: live cells. The vast majority of the cells are viable, indicating that the cluster is a living organoid. Scale bar: 100 μm.

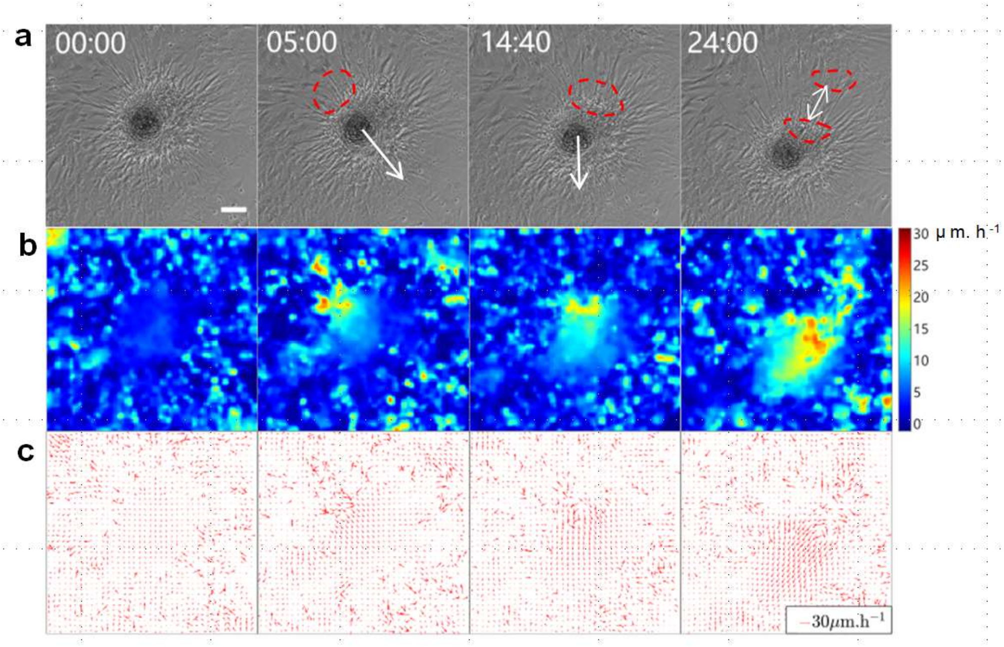


**Figure S2.** Cellular dynamics around SMC aggregates. **a:** Gradients in contractile forces around an aggregate lead to net pulling forces and aggregate displacement in the directions of the arrows. The red dashed contours demarcate zones of highly stretched cells as a result of the high pulling forces. Times shown are in hours and minutes. Scale bar: 100 µm. **b** and **c**: PIV maps demonstrating particularly elevated cellular velocities in the regions of the stretched cells.

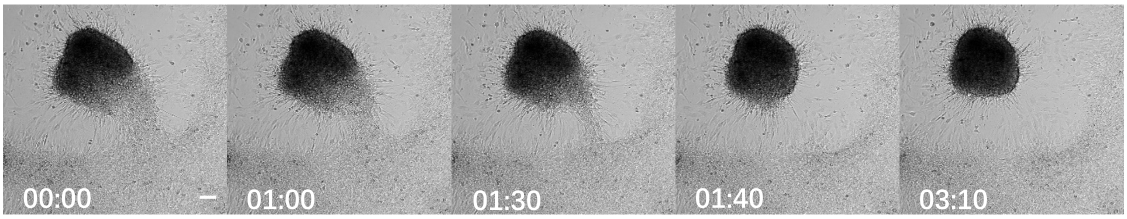


**Figure S3.** SMCs at the periphery of aggregates tear off progressively. Once the aggregate loses the last bundle of cell-substrate adhesion, it balls up into a developing cluster that eventually becomes a developed cluster with a complete contour. Scale bar: 100 µm.

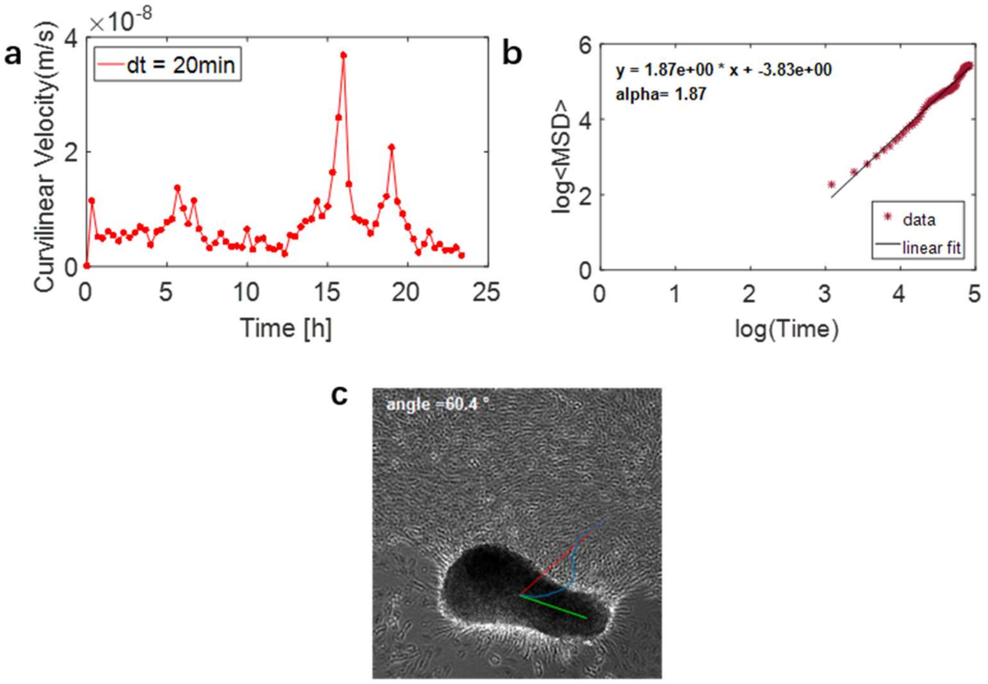


**Figure S4.** Cluster movement during the rounding-up process. **a:** Curvilinear velocity of a representative cluster as a function of time. **b:** Time evolution of the mean square displacement (MSD) for the same cluster. **c:** Cluster gets pulled by the higher SMC density on one side. Blue line denotes the cluster trajectory. Green line denotes the instantaneous direction of the cluster major axis. Red line denotes connects the initial position of the center of mass to its final position.

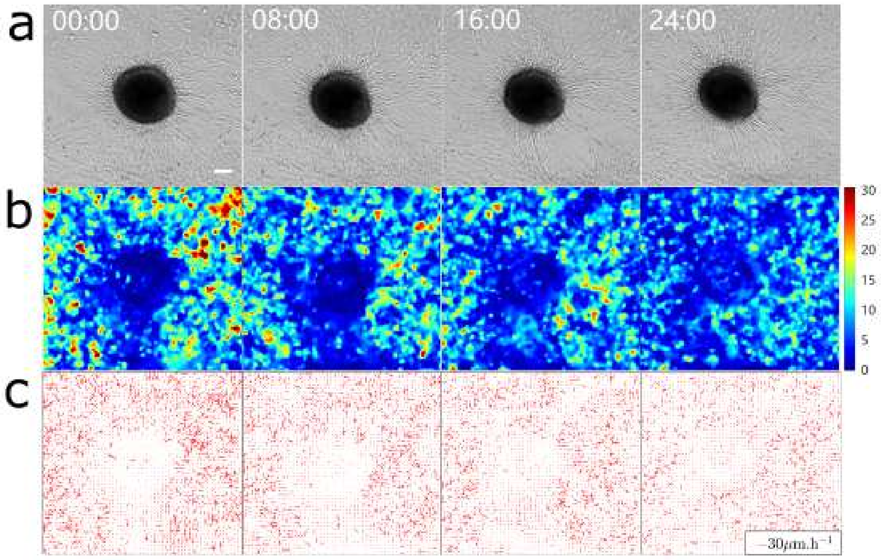


**Figure S5.** Cluster equilibrium upon stabilization. **a:** Clusters show minimal change in position after stabilization. Time points are in hours. Scale bar: 100 µm. **b** and **c:** PIV of the velocity field after stabilization. The velocities are generally low and largely symmetric around the cluster.

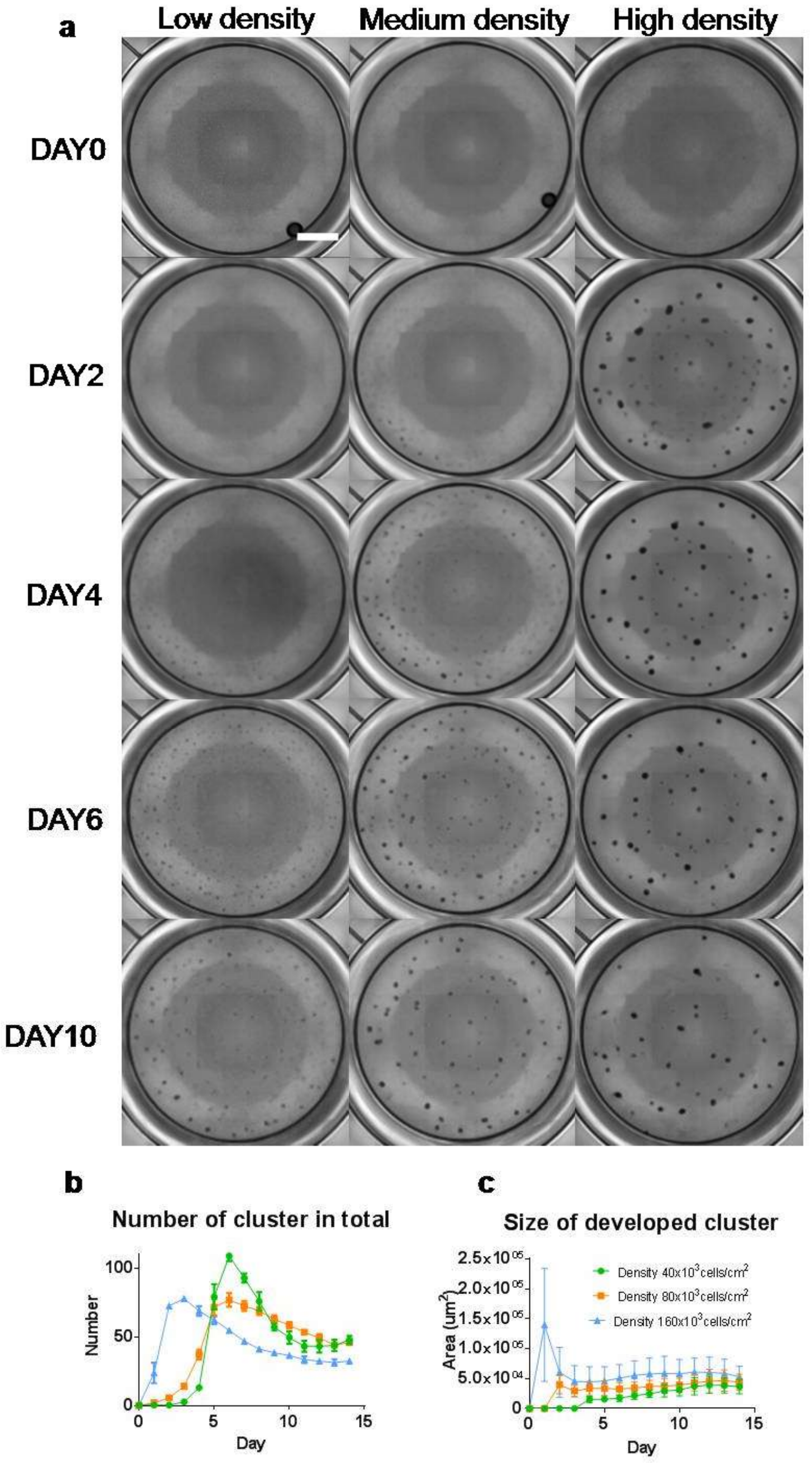


**Figure S6.** Dynamics of cluster formation at the three different cell seeding densities. **a**: Phase contrast images of cluster evolution in well-plates. Scale bar: 2 mm. **b**: Time evolution of the total number of clusters. The increase is due to cluster formation, whereas the decrease is due to cluster fusion. **c**: Time evolution of the area of developed clusters. **d** Time evolution of the number of developed cluster of different cell seeding density. The maximum number of developing clusters is reached at days 14, 9 and 4 for the low, medium and high seeding densities respectively. **e** Time evolution of the number of developing cluster of different cell seeding density. The maximum number of developing clusters is reached at days 6, 5 and 2 for the low, medium and high seeding densities respectively. **f** Time evolution of total number of cluster (sum of number of developed and developing cluster) and physical model fitting

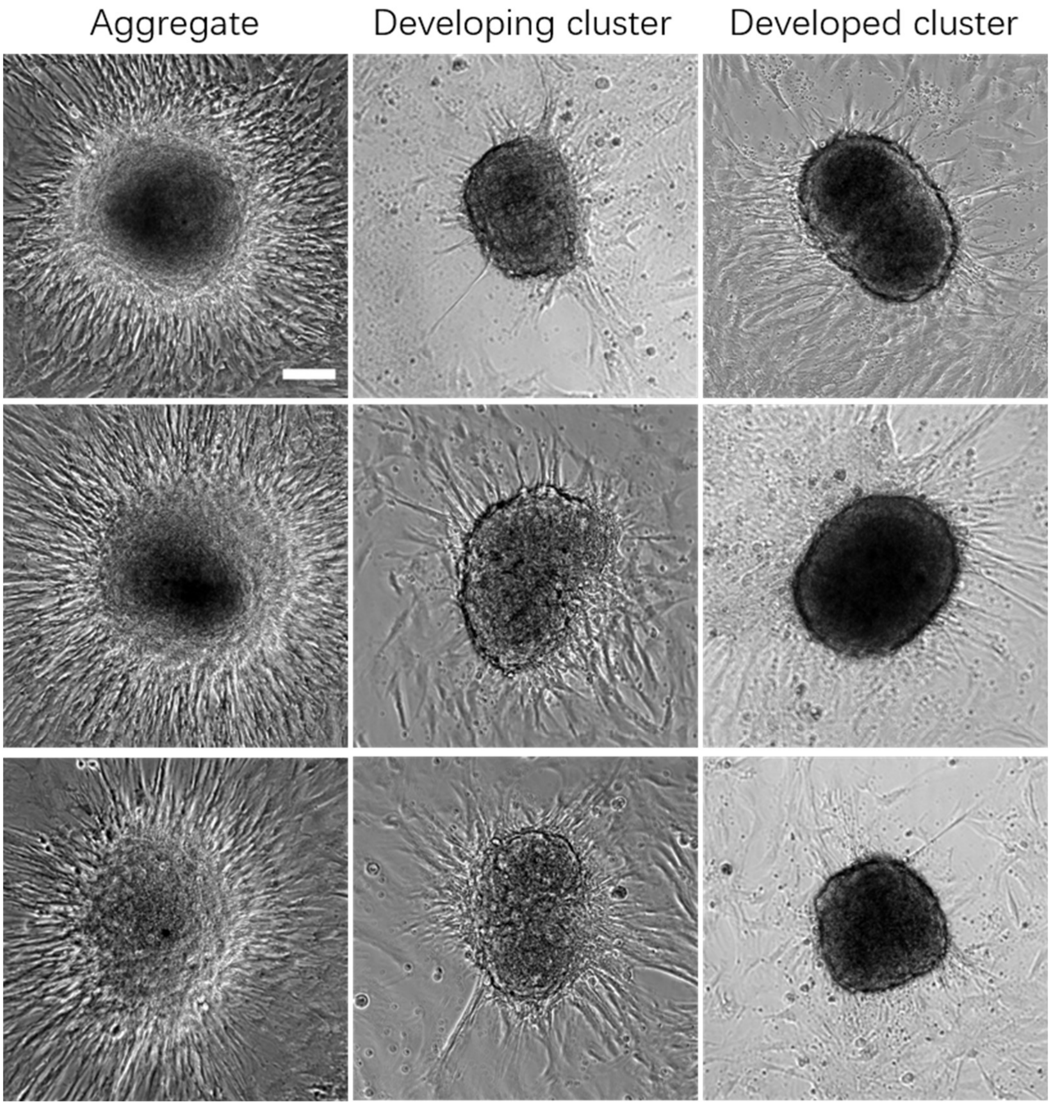


**Figure S7**. Three examples of aggregates, developing clusters and developed clusters. Aggregates, developing clusters, and developed clusters are three states seen at different times of the spontaneously arising three-dimensional SMC structures. Scale bar: 100 µm.

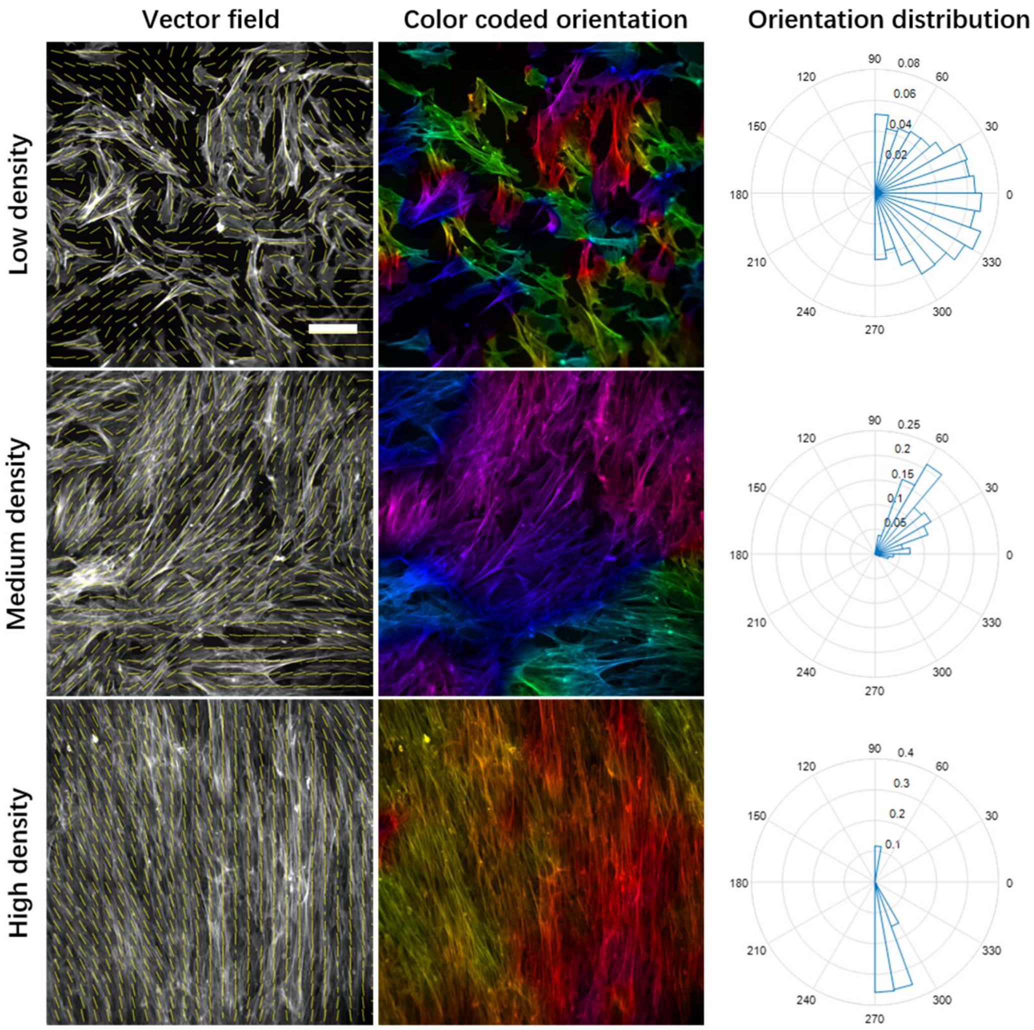


**Figure S8.** Different representations of F-actin orientation for the three different cell densities studied. Left panels: vector field of orientations. Scale bar: 100 μm. Middle panels: color-coded orientation maps. Right panels: orientation rose diagrams (polar histograms). The order parameter values for the low, medium, and high densities are as follows: $Q_{LD}$ = 0.12, $Q_{MD}$ = 0.69, $Q_{HD}$ = 0.95.

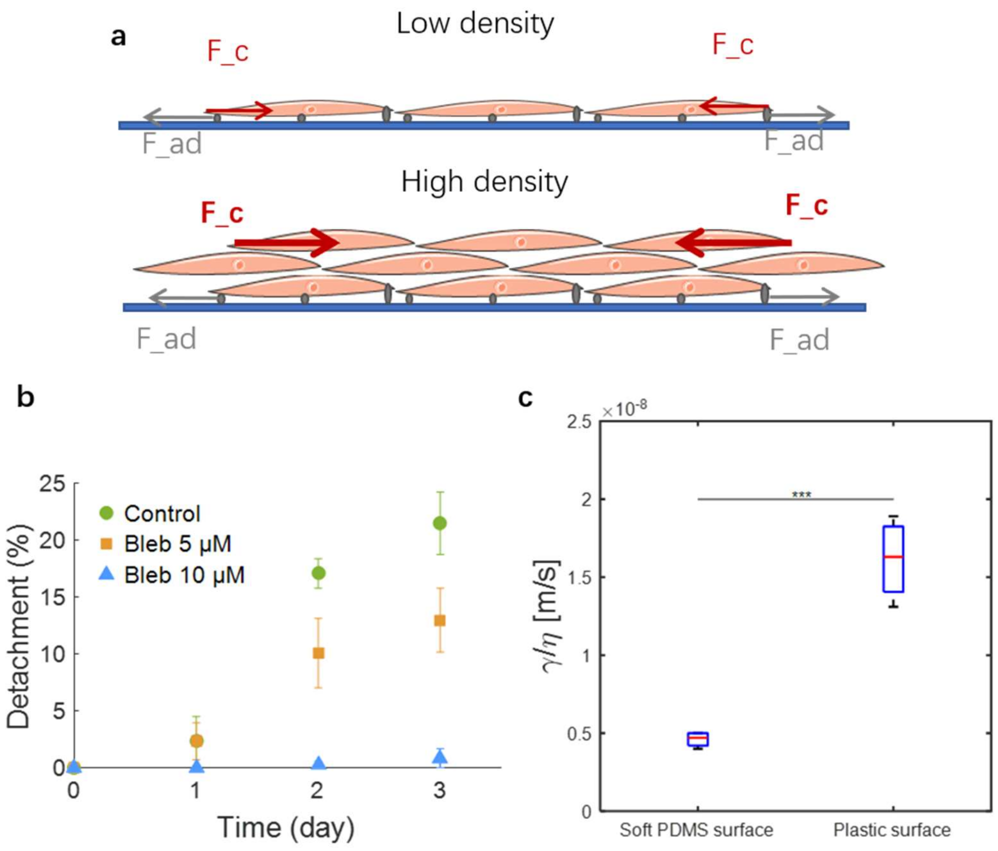


**Figure S9.** Role of contractility in SMC cluster formation. **a:** Hypothesized mechanism of cluster formation: clusters initiate when contractile forces become sufficiently large to overcome substrate adhesion forces. **b:** Contractility inhibition using blebbistatin diminishes SMC tissue detachment and cluster formation. **c:** SMCs on a soft gel where contractility is diminished exhibit a significantly smaller γ/η value, indicative of a much longer time for cluster formation than for cells on a plastic surface.

### Supplementary Notes

#### Note 1: Dynamics of event "Rounding-up"

To characterize the motion, we measured the curvilinear velocity (**Fig. S4a**) of the center of mass and the mean-square displacement (MSD) (**Fig. S4b**) of the cluster trajectory over a period of 24 h. The mean curvilinear velocity U = $7.2 \times 10^{-3}$ μm/s has the same order of magnitude as that of SMCs in a monolayer and is also comparable to that reported for the motion of aggregates of a different cell type. With an exponent α = 1.87, a value close to 2, the MSD dependence on time indicates that the SMC cluster performs a persistent walk. We followed the cluster trajectory (**Fig. S4c**) and found that the direction of persistent motion is largely orthogonal to the cluster's major axis. The cluster is often positioned between a cell-rich and a cell-free zone, and it is pulled towards the cell-rich zone by cell contraction. The contractile force exerted on a cluster is $F = kr^2U$, where r is the equivalent radius of the projected cluster area, defined above. For the measured value of $U$ and for $k \approx 2 \cdot 10^{10}$ $Pa.s.m^{-1}$, as deduced from the hole opening dynamics above, we obtain $F \approx 5$μN, which corresponds to a contractile tension $\gamma \sim \frac{F}{r} \approx 10$ mN/m, a value comparable to the surface tension reported for contractile cell aggregates[1].

**Note 2: Cluster movement during the rounding-up process**

**Curvilinear velocity**: To analyze the temporal displacement of SMC clusters, it is necessary to define an appropriate interval between successive frames for the analysis. To this end, we tested time intervals of 5, 20, and 60 min. **Figure B.1** depicts the temporal evolution of the curvilinear velocity (i.e. distance raveled per unit time) of the center of mass of a representative SMC cluster over a period of 24 h for the three time intervals, while **Error! Reference source not found.** provides the time-average curvilinear velocity and directionality for the same cluster.

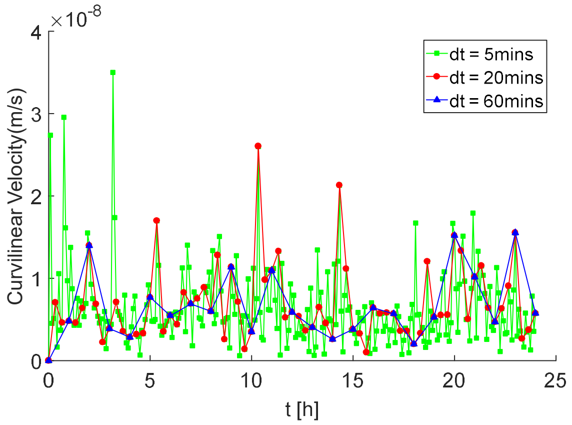


**Figure N1.** Curvilinear velocity of the centroid of a representative SMC cluster as a function of time for the three time intervals selected of 5, 20, and 60 min.

**Table N1:** Comparison of the time-average curvilinear velocity and directionality of the SMC cluster in **Fig. N1** for the three selected time intervals

| Time Interval (min) | Average curvilinear velocity (m/s) | Directionality/Persistence |
|---|---|---|
| 5 | 6.45e-9 | 0.43 |
| 20 | 6.87e-9 | 0.69 |
| 60 | 6.58e-9 | 0.80 |

These results demonstrate that the average curvilinear velocities calculated with the different time intervals are very close to one another. However, the directionality only becomes stable for time intervals of 20 min or longer. Thus, a time interval of 20 min was used in all subsequent data analysis.

**Note 3: Mathematical modeling of cluster rounding-up**

We interpret the dynamics of rounding as a phenomenon driven by cluster surface tension, arising from both cell contractility and adhesion, and resisted by cluster viscosity. Mathematically,

$$\gamma\dot{S} = -\eta \int_V \vec{\nabla}\vec{u} : \vec{\nabla}\vec{u} \mathrm{d}V, \tag{S.1}$$

where $\gamma$ is the surface tension coefficient, $\eta$ the cluster viscosity, $S$ and $V$ the cluster surface and volume, and $\vec{u}$ is the velocity field within the cluster, which we assume of the form

$$\vec{u} = \left(\frac{\rho}{R}\dot{R}, 0,0\right), \tag{S.2}$$

where $R(\theta)$ is the cluster radius and $(\rho, \theta, \varphi)$ are the spherical coordinates. We obtain:

$$\vec{\nabla}\vec{u} : \vec{\nabla}\vec{u} = \left(\frac{\dot{R}}{R}\right)^2 + \left[\frac{\partial}{\partial\theta}\left(\frac{\dot{R}}{R}\right)\right]^2. \tag{S.3}$$

We assume the cluster shape to be a prolate spheroid, of major axis *a* and minor axis *c*. We further assume the spheroid to be sufficiently close to a sphere of radius *r*. We call $\varepsilon$ the relative size of the geometric perturbation. By imposing that the deformation of the spheroid conserves a total volume equal to $\frac{4}{3}\pi r^3$, we obtain$a = r(1 - \varepsilon + \frac{2}{3}\varepsilon^2)$ and $c = r(1 + 2\varepsilon + \frac{5}{3}\varepsilon^2)$, where we neglect terms of order $\varepsilon^3$. The radius of a prolate spheroid is $R^2(\theta) = a^2\cos^2\theta + c^2\sin^2\theta$. Replacing the expressions of *a* and *c*, we obtain

$$R = r[1 + \varepsilon\, f(\theta) + \varepsilon^2\, g(\theta) + O(\varepsilon^3)], \tag{S.4}$$

where $f(\theta) = 2\sin^2\theta - \cos^2\theta$ and $g(\theta) = -\frac{1}{2}(2\sin^2\theta - \cos^2\theta)^2 + \frac{7}{6}\cos^2\theta + \frac{9}{3}\cos^2\theta$.

Replacing [4] into [3] and performing the integral in [1], we obtain the equation of the round-up dynamics,

$$\gamma\varepsilon = -\frac{56}{15}\mu\, r\, \dot{\varepsilon}. \tag{S.5}$$

To obtain the dynamical equation [4], we have supposed that the round-up occurs without changing the total aggregate volume, i.e., $\dot{r} = 0$. This is an acceptable approximation if the change in volume is small as compared to the change n shape. Solving the differential equation [4] leads to the evolution equation in the article's main text.

**Note 4: Description of "stabilization" event**

Clusters undergo changes in both shape and position during the "rounding-up" phase before eventually reaching a stable state. The cluster surface tension, arising from both cell contractility and adhesion, is balanced by cluster internal viscosity, leading to the stability in shape. The displacement of a cluster is driven by the imbalance in contraction force around it. During the "stabilization" step, the "adhering cells" around the cluster are quite homogenous, which greatly reduces the displacement of the cluster (**Fig. S5a**). We used PIV analysis to quantify the speed and direction of migrating "adhering cells" (**Fig. S5b** and **c**). The speed is generally below 30 μm/h. From the average velocity of cells at each time point, we compare the dynamics of cells from different states.

### Supplementary Movies

**Supplementary Movie 1:** Cluster formation: from homogenous seeded cell sheet to three-dimensional structure
**Supplementary Movie 2:** Dynamics of opening of hole
**Supplementary Movie 3:** Aggregate development
**Supplementary Movie 4:** Aggregate totally tear off and form a cluster
**Supplementary Movie 5:** Elongated cluster round up
**Supplementary Movie 6:** Rounded cluster stabilization
**Supplementary Movie 7:** Cluster fusion
